# Prey partitioning between sympatric canid species revealed by DNA metabarcoding

**DOI:** 10.1101/786624

**Authors:** Yue Shi, Yves Hoareau, Ellie Reese, Samuel K. Wasser

**Affiliations:** Department of Biology, University of Washington, Seattle, WA 98195, USA; School of Environmental and Forest Sciences, University of Washington, Seattle, WA 98105, USA

**Keywords:** metabarcoding, diet, wolf, coyote, fecal DNA

## Abstract

The recovery of apex predators relies on restoring the full suite of trophic interactions within the ecosystem. Diet analysis with DNA metabarcoding technology can help deliver insights into these trophic interactions with fine-grained resolution. The recovery of wolves in Washington state offers an excellent case to study the trophic cascade impacts of the apex predators on the ecosystem and explore prey partitioning between sympatric canid species. We used DNA metabarcoding technology on scats to characterize the diet composition and its spatiotemporal variations of wolves and coyotes and quantified the diet niche overlap between these two canid species in northeastern Washington. In total, 19 different prey taxa were detected. Frequency of occurrence data showed that wolves primarily preyed upon deer (*Odocoileus sp.*) (47.47%) and moose (*Alces alces*) (42.42%). Coyotes also consumed moose (30.10%) and deer (21.36%), but snowshoe hares (*Lepus americanus*) were the most common prey (61.17%) in their diet. There were significant spatial variations in the wolf diet composition (*p* = 0.001) with wolves in the Dirty Shirt pack range consuming more moose (71.43%). Coyotes showed significant spatial and temporal dietary variations (season: *p* = 0.037; pack: *p* = 0.003; pack:season *p* = 0.043). Our data suggested that coyotes use ungulate carrion subsidies from wolves as food resources. DNA metabarcoding with fecal DNA provides an excellent noninvasive tool to characterize diet profile at the fine-grained level and can be applied to other carnivore species to help understand the impacts of recovery of apex predators on the local ecosystems.

## Introduction

Apex predators are primarily known for their elevated position on the trophic ladder with impacts that cascade throughout their ecosystem (Wallach, Izhaki, Toms, Ripple, & Shanas, 2015). The widespread decline in apex predators due to human hunting and habitat fragmentation has been observed in many systems (Estes et al., 2011). The reestablishment of large predators and their ecological effects is fundamental for wildlife management, and relies on restoring the full suite of ecological interactions within the ecosystem, including predator-prey and predator-predator interactions (Stier et al., 2016). Scientific uncertainty about the ecological interactions can hinder the progress towards resolving the new conservation and legal challenges introduced by apex predator recovery (Marshall, Stier, Samhouri, Kelly, & Ward, 2015). Therefore, it is essential to track the dietary changes of predators to inform conservation management decisions (Marshall et al., 2015). Diet analysis with the newly developed DNA metabarcoding technology can help deliver valuable insights into these trophic interactions and the mechanism of species coexistence with the fine-grained resolution.

Conventional methods of diet analysis have relied on macro- or microscopic morphological identification of food remains in scats, such as hair (Carrera et al., 2008; Gable, Windels, Bruggink, & Barber-Meyer, 2018; Wasser, Keim, Taper, & Lele, 2011) and hard-parts (e.g. bones, hooves and teeth) (Drouilly, Nattrass, & O’Riain, 2017; Nelson, Cherry, Howze, Warren, & Mike, 2015). Such methods are very labor intensive and require reliable reference collection of prey parts as well as research expertise in identifying species from masticated, semi-digested food remains (Pompanon et al., 2011). These methods also tend to underestimate the proportion of prey species whose remains are not found (completely digested or not consumed at all) (Deagle, Kirkwood, & Jarman, 2009). Earlier molecular attempts use a conventional genotyping methodology, involving multiplex PCR followed by fragment size determination with capillary electrophoresis and known sized allelic ladders (Morello, Braglia, Gavazzi, Gianì, & Breviario, 2019). Though very affordable, this method does not account for any sequence-based differences between fragments of the same size. DNA metabarcoding offers a promising alternative, whereby customized universal primer pairs amplify a standardized DNA region, which are sequenced and compared to a reference database for taxonomic identification (Modave, MacDonald, & Sarre, 2017; Taberlet, Coissac, Pompanon, Brochmann, & Willerslev, 2012). Coupled with next-generation sequencing (NGS), DNA metabarcoding technology can sequence many samples in a high-throughput and cost-effective fashion and reveal the entire taxonomic composition of thousands of samples simultaneously (Pompanon et al., 2011). This new molecular approach has been successfully applied for the diet analyses of various species, including carnivores (Berry et al., 2017; Smith, Thomas, Levi, Wang, & Wilmers, 2018), omnivores (De Barba et al., 2013; Robeson et al., 2017), herbivores (Kartzinel et al., 2015), small mammals (Buglione et al., 2018), lizards (Moreno-Rueda, Melero, Reguera, Zamora-Camacho, & Álvarez-Benito, 2017), birds (Sullins et al., 2018) and invertebrates (Hawlitschek, Fernández-González, Balmori-de la Puente, & Castresana, 2018; Kamenova, Bretagnolle, Plantegenest, & Canard, 2018).

Gray wolves (*Canis lupus*) were extirpated from Washington state by the 1930s as ranching and farming activities expanded (Wiles, Allen, & Hayes, 2011). Coyotes (*Canis latrans*) increased in range and abundance over the same period (Gallagher et al., 2019; Hody & Kays, 2018). Since 2008, wolves have begun reestablishing territories in Washington through natural dispersal from adjacent states and provinces such as Idaho, Montana, Oregon, and British Columbia (Wiles et al., 2011) after being absent from the state for over 80 years. The recovery of wolves in Washington state offers an excellent case to study the trophic cascade impacts of the apex predators on the ecosystem and explore prey partitioning between sympatric canid species.

As apex predators, wolves are adapted to prey on large ungulates (Gable et al., 2018; Wasser et al., 2011). Wolves are also opportunists and use small prey as seasonal food sources when abundant (Latham, Latham, Mccutchen, & Boutin, 2011). Coyotes are mesopredators and generally viewed as opportunistic generalists (Kilgo, Vukovich, Ray, Shaw, & Ruth, 2014). The majority of the coyote diet consists of small mammals, but coyotes can also prey on ungulate calves (Chitwood et al., 2015; Kilgo, Ray, Vukovich, Goode, & Ruth, 2012; Nelson et al., 2015), and vulnerable ungulate adults (Benson, Loveless, Rutledge, & Patterson, 2017; Patterson & Messier, 2000). Birds represent a nonnegligible proportion of the coyote diet (Smith et al., 2018). In addition, carrion subsidies from wolves is also a highly valued food resource for coyotes (Sivy, Pozzanghera, Colson, Mumma, & Prugh, 2017). Using traditional methods, these previous studies generally present prey species as three groups: ungulates, small mammals, and birds. All these groups encompass enormous taxonomic diversity, yet very few studies have evaluated resource partitioning of these sympatric canid species at a fine-grained taxonomic level.

In order to recover apex predators, it is critical to consider the ecological roles that these top predators play in the ecosystem, rather than focusing only on their demography (Ripple, Wirsing, Beschta, & Buskirk, 2011). Here, we use DNA metabarcoding on scats to construct high-resolution diet profiles of sympatric wolves and coyotes in northeastern Washington state, USA, with a focus on the vertebrate component of their diets. The purpose of the study is to: 1) evaluate the effectiveness of DNA metabarcoding approach for the canid diet analysis; 2) assess the necessity of applying predator-specific blocking primers to construct a full profile of prey composition. Predator DNA could swamp prey DNA during amplification because predator DNA is much more abundant. The predator-specific blocking primer was designed in a way that its 3’ end was modified by replacing the 3’ hydroxyl group with a spacer-C3-CPG to prevent polymerase extension (Vestheim & Jarman, 2008); 3) examine spatiotemporal variation in the diet profiles of wolves and coyotes as both pack-specific prey abundance and hunting-ability could affect prey consumption rates. We believe it is crucial to develop a holistic picture of what dietary options these sympatric canid predators exploit in different ecological contexts.

## Materials and Methods

### Ethics Statement

Currently, wolves in the western two-thirds of Washington are listed as endangered under federal law. Wolves in the eastern third of the state have been removed from the federal listing. However, all wolves in Washington are listed as endangered under state law (Wiles et al., 2011). Three recovery regions have been delineated in Washington: (1) Eastern Washington, (2) Northern Cascades, and (3) Southern Cascades and Northwest Coast. Our study area is in the Eastern Washington recovery region. Coyotes are not protected under the Endangered Species Act anywhere in the contiguous United States. Fecal samples were collected using the detection dogs from the Conservation Canine Program at the University of Washington under IACUC protocol #2850-08.

### Sample Collection

Fecal samples were collected in the spring season (April and May in 2015 and 2017) and fall season (October and November in 2015 and 2016) in three wolf pack ranges: Smackout, Dirty Shirt and Goodman Meadows in Pend Oreille and Stevens counties, Washington (Figure 1). Upon collection, fecal samples were stored at −20°C until DNA extraction. Samples used in this study were subsampled from a larger ongoing project (wolf: 647 fecal samples; coyote: 1893 fecal samples) (Supplementary Figure 1). Subsampling was based on relative kernel density, which was calculated using *stat_density2d* function in *ggplot2* in R (Figure 1). We selected samples from the areas with highest and lowest estimated wolf and coyote densities in each of the three wolf pack ranges.

**Figure 1.**
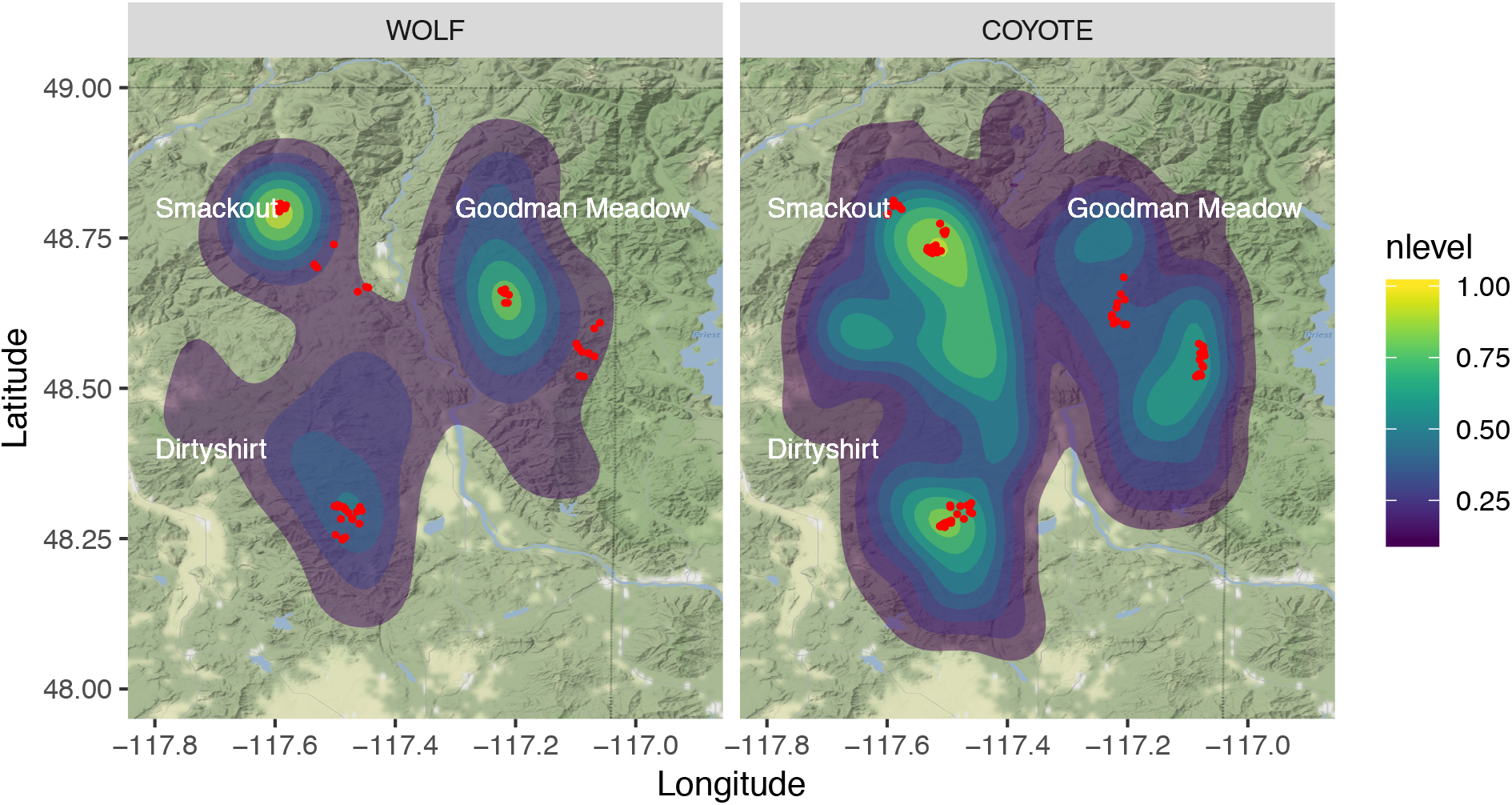
Sample selection based on relative kernel density (nlevel) map. Samples used in this study were selected from an ongoing project with 647 wolf fecal samples and 1893 coyote fecal samples. Kernel density was calculated using function *stat_density2d* in R *ggplot2*. Note: The color gradient indicates the relative density ranges from 0 to 1.

### DNA Extraction and Species Identification

Fecal DNA was extracted using the swabbing method described previously (Wasser et al., 2011) with Qiagen DNeasy 96 Blood and Tissue Kit (Qiagen Inc.,). Multiple interior surfaces of each fecal sample were swabbed and extracted in duplicate. Extraction duplicates were pooled before PCR. One extraction blank was processed along with every 45 fecal samples. Predator species identification was conducted by targeting the partial mitochondrial control region, which was amplified in duplicate using 5’6-FAM-labeled LTPROBB13 (5’-CCACTATTAACACCCAAAGC-3’) and unlabeled HSF21 (5’ - GTACATGCTTATATGCATGGG-3’) primers, following the instruction of Qiagen Multiplex PCR kit with annealing temperature at 60°C. Fragment analysis on a 3730 Genetic Analyzer (Applied Biosystems) was used to identify a fragment size of 170 bp unique to wolves and domestic dogs, and a fragment size of 165bp unique to coyotes. Samples identified as wolf/dog were further delineated by sequencing a 208bp cytochrome b fragment (Reese et al. 2019, in prep).

### Library Preparation and Sequencing

We conducted in-silico analyses using ecoPrimer (Riaz et al., 2011) and ecoPCR (Ficetola et al., 2010) before conducting the experiments to ensure that all target species could be successfully amplified with 12S V05F/R primers (Riaz et al., 2011). We followed the two-step library preparation protocol modified from Illumina’s 16S Metagenomic Sequencing Library Preparation (CT #: 15044223 Rev. B). Our protocol used customized library indices, which incorporated two distinct 10-bp index sequences on each end of the fragment. We first amplified the 12S rRNA gene V5 region using the 12SV05F/R primer pair with the addition of the overhang sequence to allow the subsequent annealing of index primers. Amplicon libraries were then amplified with index primers, which incorporated indices and Illumina sequencing adapters. We also tested the efficiency of a 10-fold excess of wolf-specific blocking oligonucleotide (via personal communication with Dr. Shehzad Wasim, University of Veterinary & Animal Sciences, Lahore, Pakistan). Since wolves and coyotes share the same sequence in the target 12S V5 region, we used the same blocking primer for both wolf and coyote samples. In total, we conducted three PCR replicates without predator-specific blocking primer, and three PCR replicates with predator-specific blocking primer. All PCR reactions were performed using the Qiagen Multiplex PCR kit along with PCR negative controls (5 PCR negative controls per 96-well plate). Amplification products were purified with 1.8X SPRI bead solution after each PCR step (Rohland & Reich, 2012). See Supplementary File 1 for a detailed description of the library preparation protocol and primer sequences.

Libraries were quantified using the Qubit dsDNA HS (High Sensitivity) Assay Kit (Invitrogen) and checked for integrity using Agilent 2200 TapeStation High Sensitivity DNA 1000 Kit (Santa Clara, CA). Successful libraries were pooled with an equimolar concentration of 2.5 nM, and the final library pool was further diluted to 2 nM. For specific methods about library QC and pooling, see Supplementary File 2. Sequencing was performed on the MiSeq platform using MiSeq Reagent kit v3 and 150 bp pair-end read length configuration. Library pool was loaded at a loading concentration of 8 pM with 25% PhiX control V3 spike-in to improve the quality of low-diversity libraries.

### Sequence Analysis and Taxonomic Assignment

MiSeq automatically separated all reads by samples during the post-run process via recognized indices. Filtering of the sequences and taxonomic inference of molecular operational taxonomic units (MOTUs) was performed using the OBITools package (Boyer et al., 2015). The following steps were performed: 1) merge paired reads with *illuminapairedend* command and filter out reads with alignment score less than 200; 2) add “sample” attribute to all reads with *obiannotate* command; 3) concatenate all reads and dereplicate globally using *obiuniq* command while keeping sample attribute; 4) remove reads that are shorter than 80 bp and less than 400 copies with *obigrep* command; 5) remove PCR and sequencing errors with *obiclean* command; 6) assign taxonomy for each MOTU to the species or genus level using the *blastn* with e-value < 1 × 10^−20^ and a minimum identity of 0.98. Taxonomic assignment was restricted to the local species in Washington state, compiled from three sources of information: 1) Mammal collection at the Burke Museum, University of Washington (https://www.burkemuseum.org/research-and-collections/mammalogy/collections/mamwash/); 2) BirdWeb (http://www.birdweb.org/birdweb/); 3) Washington NatureMapping Program (http://naturemappingfoundation.org/natmap/maps/wa/).

Further filtering on the MOTU table was conducted in R using the following steps: 1) remove any MOTU whose relative frequency across the entire dataset was found to be maximum in either extraction or PCR negative controls; 2) subtract the maximum abundance in extraction or PCR negative controls of the remaining MOTUs from their abundance in each sample replicate; 3) suppress the read count of any MOTU in a sample replicate to zero when its relative abundance (abundance in a sample replicate / total abundance across the entire dataset) is below 0.03%. The sequence read count data were converted into a MOTU table with presence/absence data. The reliability of extracting quantitative information with relative read abundance (RRA) is questionable because of variations in tissue cell density, gene copy, survival rates of tissue/DNA during digestion, variance in fragment size among different food items, and, more importantly, PCR bias due to primer-template mismatches (Piñol, Mir, Gomez-Polo, & Agustí, 2014; Pompanon et al., 2011). Therefore, we only used presence/absence data for the following analyses. However, we also generated a diet profile for each predator with RRA data for the purpose of comparison. A specific prey item in a given sample was only considered as present if it was detected in at least 2 replicates.

### Statistical Analyses

Frequency of occurrence (FOO) of a prey species was defined as the proportion of scats found to contain that prey item. To test if there was any significant interspecific and spatiotemporal variation in diet composition, we conducted permutational multivariate analysis of variance (perMANOVA) using the *adonis2* function in vegan package with the *jaccard* distance matrix and 999 permutations. Pairwise comparisons were conducted using the similarity percentage (SIMPER) test implemented in *vegan* R package with 999 permutations. Significance levels were adjusted with sequential Bonferroni correction for multiple comparisons. The SIMPER function also reports the contribution of each prey item to the overall diet dissimilarity and displays the most important prey item for each pair of comparison. These important species contribute at least to 70% of the differences between each pair of comparison. The rest of species are considered to have minor contributions. We used Pianka’s adaptation of the niche overlap (*O* metric) (Pianka, 1973) to determine dietary overlap between wolves and coyotes in different seasons or wolf pack ranges. The *O* metric ranges from 0 (no overlap) to 1 (complete overlap).

## Results

### Prey Species Detected in the Study Area

We selected 99 genetically confirmed wolf scats and 103 genetically confirmed coyote scats. In total, 19 different prey species were identified, including 5 ungulate species (deer *Odocoileus sp.*, moose *Alces alces*, elk *Cervus canadensis*, domestic cow *Bos taurus*, and pig *Sus scrofa*), 10 small mammals (snowshoe hare *Lepus americanus*, deer mouse *Peromyscus maniculatus*, meadow vole *Microtus pennsylvanicus*, red squirrel *Tamiasciurus hudsonicus*, ground squirrel *Spermophilus sp.*, red-backed vole *Myodes sp.*, flying squirrel *Glaucomys sp.*, rabbit *Oryctolagus cuniculus*, muskrat *Ondatra zibethicus*, and chipmunk *Tamias sp*.) and 4 bird species (ruffed grouse *Bonasa umbellus*, wild turkey *Meleagris gallopavo*, common starling *Sturnus vulgaris*, and spruce grouse *Dendragapus canadensis*).

### Effects of Predator-Specific Blocking Primer

The use of a predator-specific blocking primer increased the proportion of prey sequences from 26.40% to 65.97% (Figure 2). However, all prey species were detected regardless of whether the predator-specific blocking primer was added. The blocking primer increased the detection of 9 prey species, while reducing the detection of chipmunk, domestic cow, muskrat, pig and snowshoe hare (Supplementary Table 1). The detection of common starling, ground squirrel, rabbit, spruce grouse and wild turkey remained the same (Supplementary Table 1). Chipmunk was not detected with addition of the blocking primer. For the subsequent analyses, we combined the data from all 6 PCR replicates of each sample. A specific prey item in a given sample was only considered present if it occurred in at least 2 out of 6 replicates.

**Figure 2.**
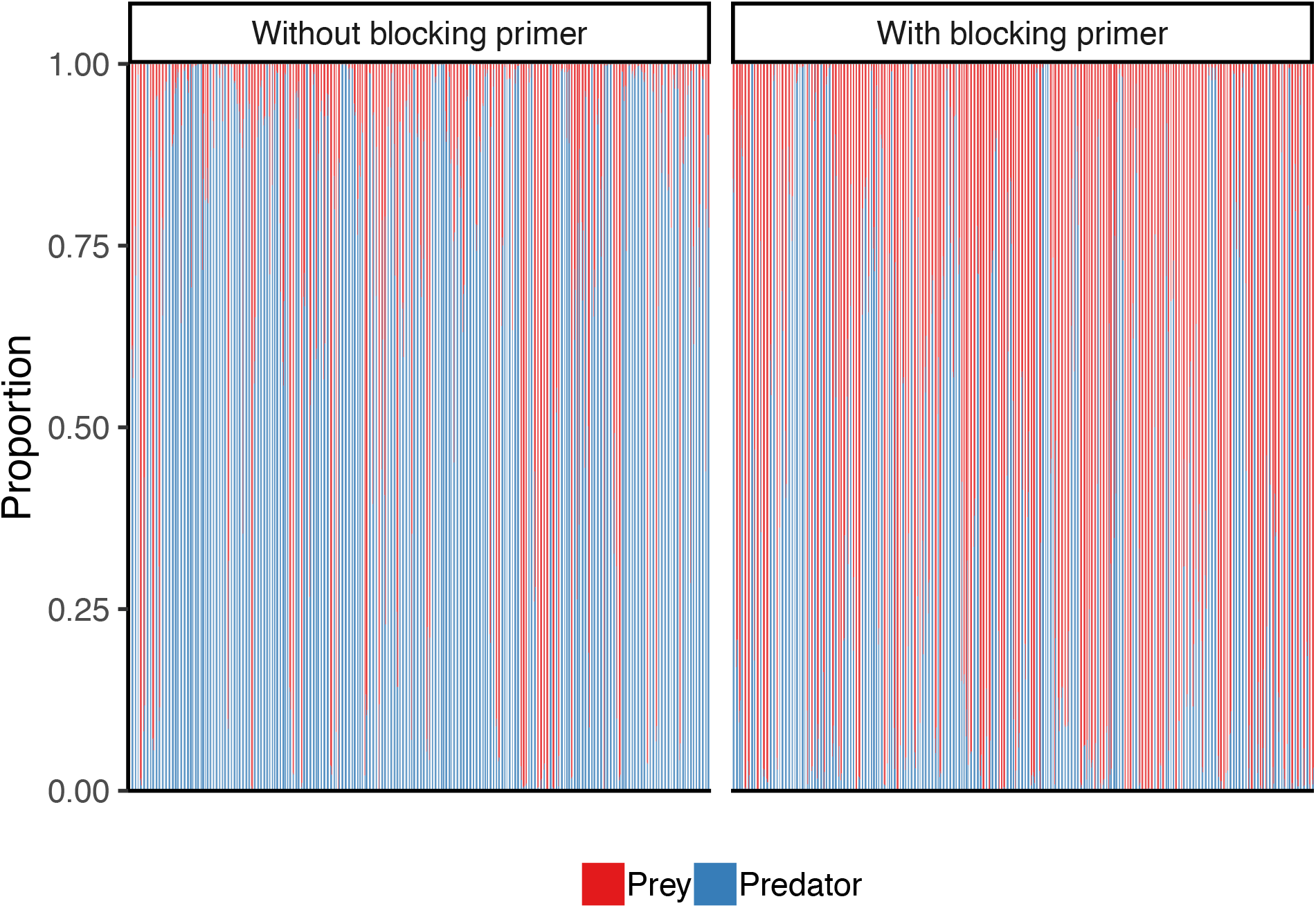
Effects of the predator-specific blocking primer on the proportion of reads mapped to the predators and preys.

### Interspecific Differences in Diet Profiles

The diet compositions of wolves and coyotes were significantly different (*p* = 0.001) (Figure 3). Deer and moose were the two most frequent prey species (47.47% and 42.42% respectively) in the wolf diet, followed by elk (17.17%) and domestic cow (16.16%). Domestic cow was detected in 16 out of 99 wolf scats (Supplementary Table 3). By contrast, snowshoe hares were the most common prey species (61.17%) in the coyote diet, followed by moose (30.10%) and deer (21.36%) (Figure 3). Domestic cow was only detected in 2 out of 103 coyote scats (Supplementary Table 3). Small mammals and birds were rarely detected in the wolf diet, whereas they occurred relatively more common in the coyote diet (Figure 4). In total, we found 11 prey species in the wolf diet and 18 prey species in the coyote diet. On average, there were 1.48 prey species per wolf scat, and 1.79 prey species per coyote scat. The SIMPER test showed that snowshoe hares (*p* = 0.001) and deer (*p* = 0.002) were the most influential prey species, significantly contributing to the interspecific dietary differences. Domestic cow, deer mouse, ground squirrel, meadow vole and red-backed vole also significantly contributed (*p* < 0.05) to the interspecific dietary differences, through their contributions were minor. The diet profiles of wolves and coyotes generated with RRA data was similar to those with FOO data (Supplementary Figure 2), except that each prey species had lower proportion with RRA data. For example, the average read relative abundance of domestic cow was 3.73% with RRA data, whereas the its FOO was 16.16%.

**Figure 3.**
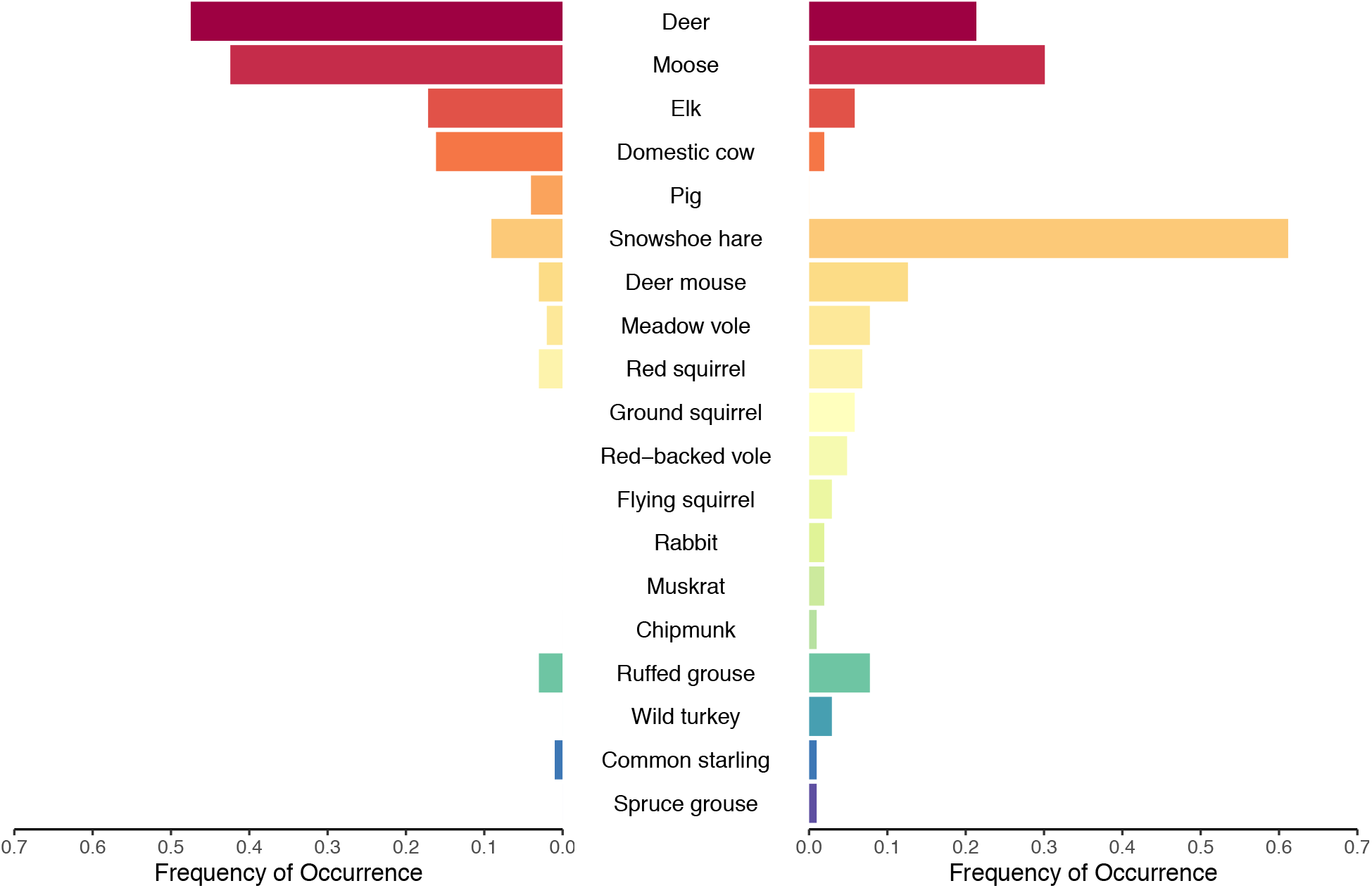
Diet profile of wolves (*N* = 99) and coyotes (*N* = 103) using the frequency of occurrence of 19 prey species. There was a significant difference in the diet profile between wolves and coyotes (*p* = 0.001).

**Figure 4.**
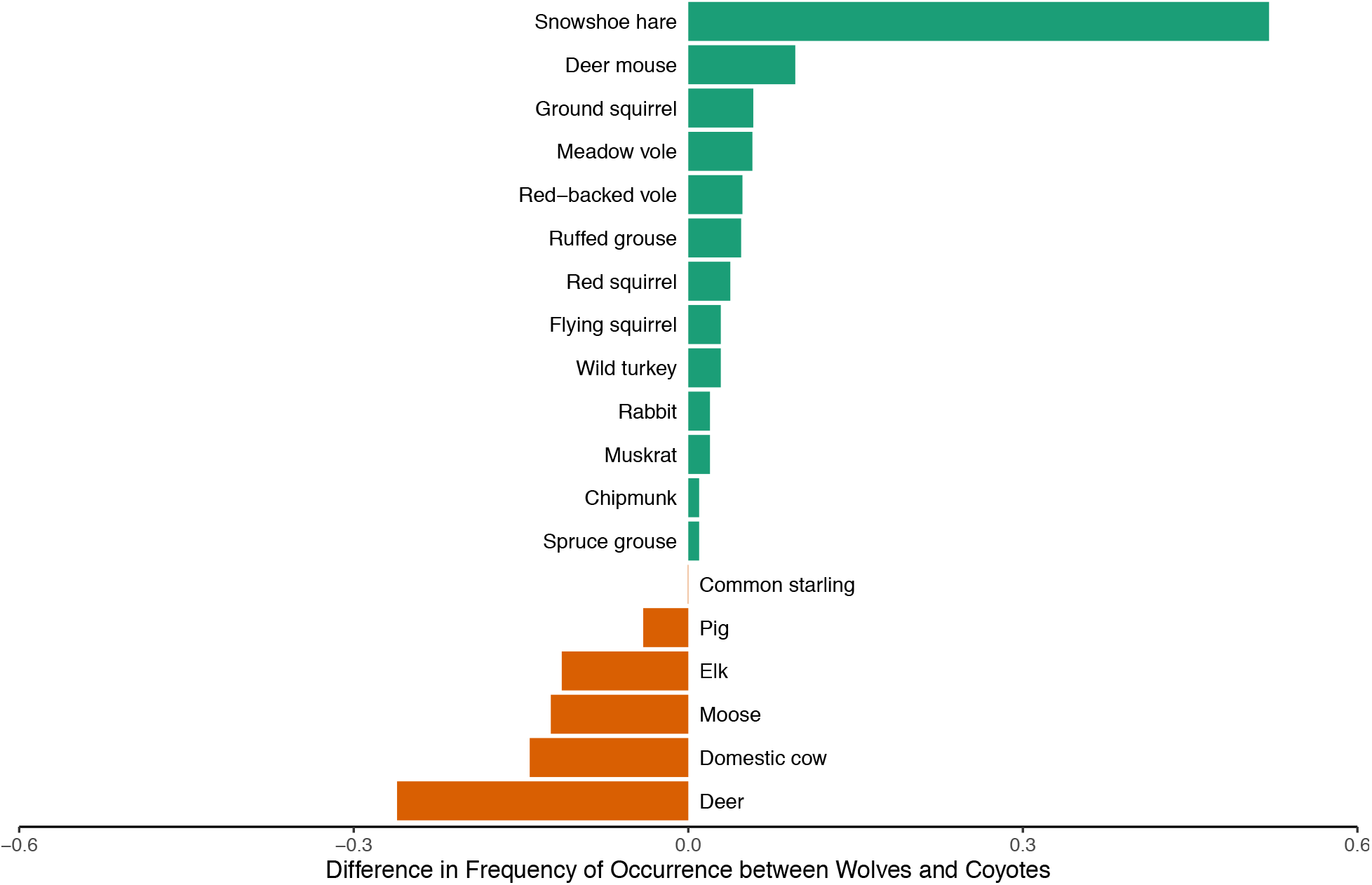
Differences in diet profile of wolves (*N* = 99) and coyotes (*N* = 103) using the frequency of occurrence of 19 prey species. Difference in the frequency of occurrence (FOO) of any prey species was calculated as FOO in wolf - FOO in coyote.

### Spatiotemporal Variations in the Diet Profile of Wolves

The dietary differences among wolf pack ranges were significant (*p* = 0.001). As there was no significant seasonal difference in the wolf diet (*p* = 0.448) or significant interaction between seasons and wolf pack ranges (*p* = 0.095), following analyses focused only on the spatial variation in the wolf diet (Figure 5). The FOO of moose was the highest (71.43%) in the Dirty Shirt range, followed by Smackout (38.24%) and Goodman Meadows (24.32%). According to the SIMPER test, moose consumption in the Dirty Shirt range was significantly higher than that in Goodman Meadows (*p* = 0.001). The FOO of deer was highest in the Smackout range (58.82%), followed by Goodman Meadows (51.35%) and Dirty Shirt (28.57%). The FOO of elk was the highest in the Goodman Meadows range (29.73%), whereas its FOO was 11.76% and 7.14% in the Smackout and Dirty Shirt ranges, respectively. Common starling was only found in the Goodman Meadows range with a FOO of 2.70%. Meadows vole and ruffed grouse were only found in the Smackout and Dirty Shirt ranges (Figure 5).

**Figure 5.**
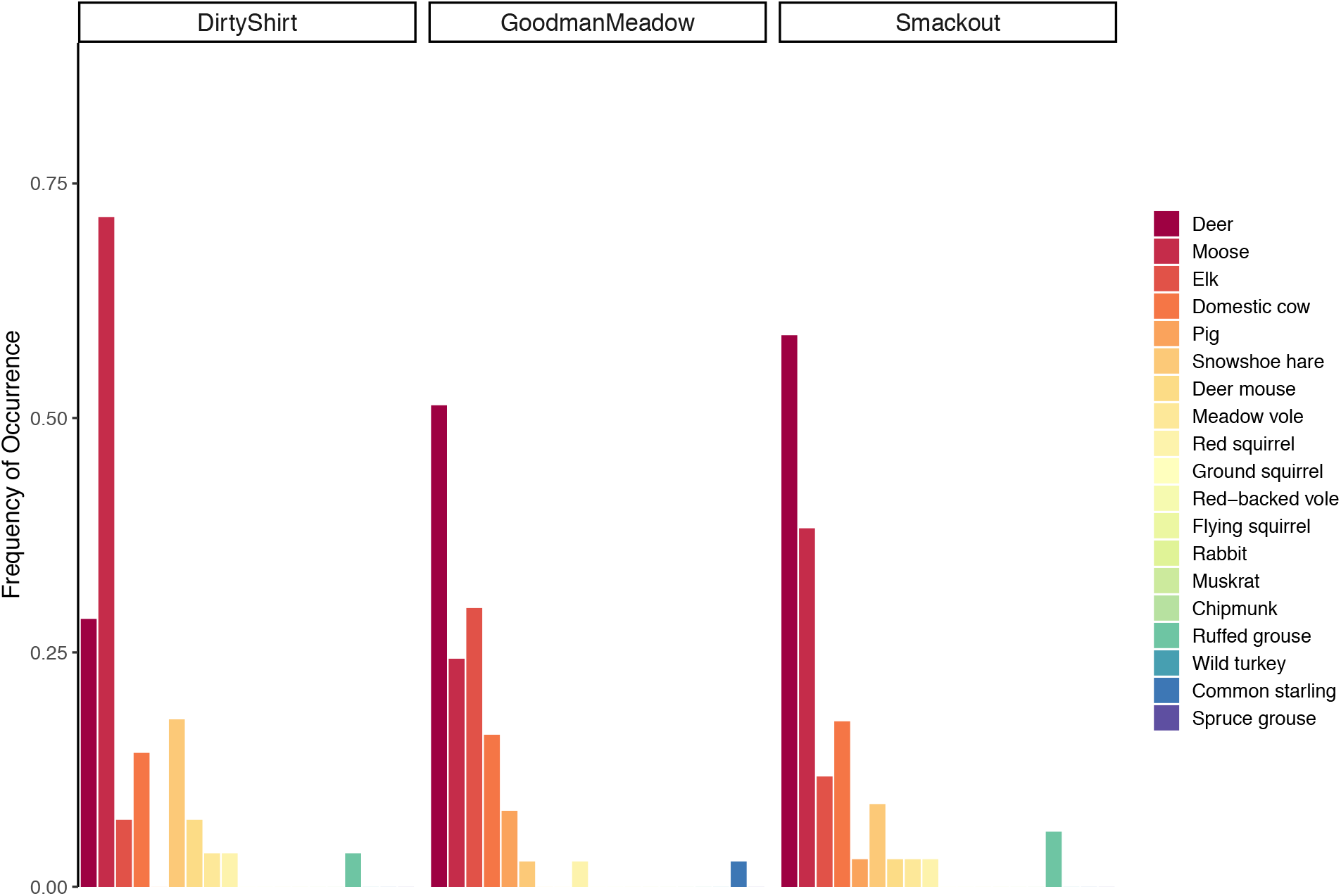
Significant spatial differences in the wolf diet profiles among three wolf pack ranges, using the frequency of occurrence of 19 prey species (*p* = 0.001).

### Spatiotemporal Variations in the Diet Profile of Coyotes

There were significant spatiotemporal variations in the coyote diet (season: *p* = 0.037; pack: *p* = 0.003) (Figure 6). Since there was a significant interaction between seasons and wolf pack ranges (*p* = 0.043), we investigated the spatial dietary changes in coyotes in each season separately. In the spring, the FOO of moose in the Dirty Shirt range was highest (66.67%) among three wolf pack ranges, whereas the FOO of moose in the Smackout range was the lowest (12.50%). The difference in FOO of moose between Dirty Shirt and Smackout was significant (*p* = 0.001). In the fall, deer was not detected in the coyote diet in the Goodman Meadows range, whereas its FOO was highest in Smackout (60.00%). Muskrat was only detected in the Goodman Meadows range (*p* = 0.014) and wild turkey was detected in Dirty Shirt and Smackout ranges but not in Goodman Meadows (*p* = 0.015). We also investigated the seasonal dietary changes of coyotes in each wolf pack range separately. In the Dirty Shirt range, moose consumption was higher in the spring (66.67%) than in the fall (15.38%), though the difference was not significant after correcting for multiple comparisons (*p* = 0.021, adjusted alpha=0.017). In the Goodman Meadows range, muskrat consumption was only detected in the fall (*p* = 0.011). In the Smackout range, deer consumption was significantly higher in the fall (60.00%) than that in the spring (20.83%) (*p* = 0.012).

**Figure 6.**
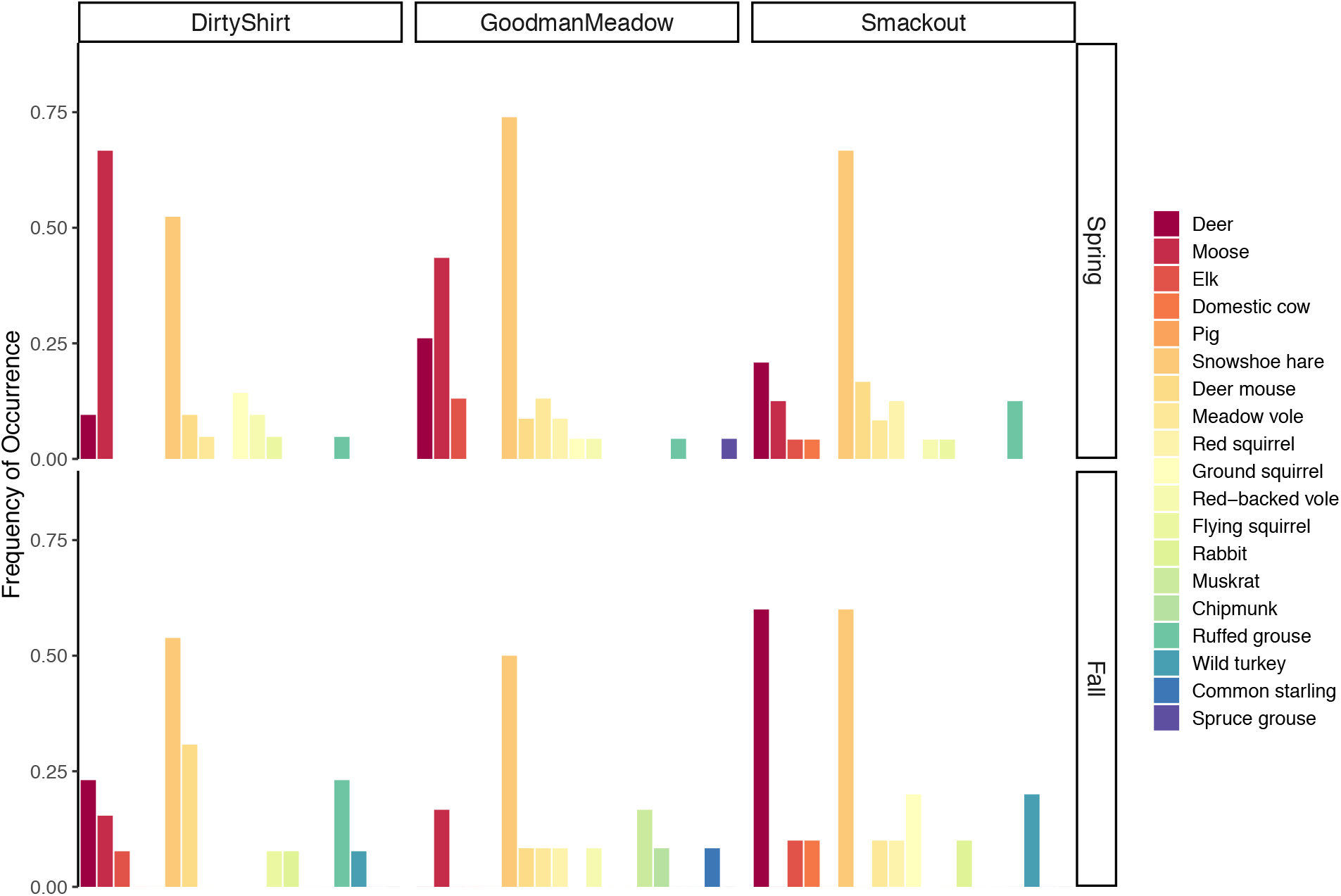
Spatiotemporal variations in the diet profile of coyotes, using the frequency of occurrence of 19 prey species. Both the main terms and the interaction term were significant based on perMANOVA analysis (season *p* = 0.037; Pack: *p* = 0.003; pack:season *p* = 0.043).

### Spatiotemporal Variations in Diet Overlap between Wolves and Coyotes

There was large variation in dietary overlap (*O*) between wolves and coyotes, ranging from nearly no overlap (0.08) to 0.74. Dietary overlap was least in the Goodman Meadows range in the fall, at 0.08. The most substantial dietary overlap (*O*) between wolves and coyotes was found in the Smackout range in the fall (0.74), followed by the Dirty Shirt range in the Spring (0.70) (Table 1).

**Table 1.** Dietary overlap between wolves and coyotes in different seasons or wolf pack ranges. Dietary overlap was determined with Pianka’s adaptation of the niche overlap (*O* metric), ranging from 0 (no overlap) to 1 (complete overlap). Sample size (N) for each predator in each season or wolf pack range was also given.

|  | Dirty Shirt |  | Goodman Meadows |  | Smackout |  |
| --- | --- | --- | --- | --- | --- | --- |
|  | Spring | Fall | Spring | Fall | Spring | Fall |
| N (Wolf) | 7 | 21 | 13 | 24 | 24 | 10 |
| N (Coyote) | 21 | 13 | 23 | 12 | 24 | 10 |
| *O* metric | 0.70 | 0.56 | 0.58 | 0.08 | 0.41 | 0.74 |

## Discussion

In this study, we characterized the high-resolution diet profiles of sympatric wolves and coyotes in northeastern Washington, and also revealed their dietary spatiotemporal variations. We demonstrated that DNA metabarcoding was a successful and effective molecular approach to track dietary changes of predators and help inform conservation management decisions.

### Prey Partitioning between Sympatric Canids in Northeastern Washington

The results with FOO data showed that wolves primarily preyed on deer (47.47%) and moose (42.42%) in northeastern Washington. This result is consistent with kill site analyses conducted by Washington Department of Fish and Wildlife (WDFW) (Kertson, 2018). Wolves hunted moose across much of the boreal forest of North America (Latham, Latham, Knopff, Hebblewhite, & Boutin, 2013). We couldn’t determine the specific deer species with 12S V5 sequence. In northeastern Washington, there are two deer species, mule deer (*Odocoileus hemionus*) and white-tailed deer (*Odocoileus virginianus*). White-tailed deer is the predominant deer species as there is limited mule deer habitat in the study area (Washington Department of Fish and Wildlife, 2017). Snowshoe hares (61.17%) were the most common prey in the coyote diet, which is consistent with previous studies (Latham et al., 2011; Smith et al., 2018). The high snowshoe hare consumption rate by coyotes has significant impacts on the conservation management of Canada lynx (*Lynx canadensis*). Lynx are usually considered as specialist on snowshoe hares, whereas coyotes are often considered as generalist. The roughly 10 - year population cycle of snowshoe hare is among the most well-known examples of cyclic population dynamics (Stenseth, Falck, Bjornstad, & Krebs, 1997). Coyote and lynx both mostly feed on snowshoe share except during its cyclic lows (O’Donoghue et al., 1998). In 2000, Canada lynx was listed as threatened under the US Endangered Species Act (ESA) and in 2016, Canada lynx was listed as endangered in Washington state. In the absence of wolves, coyotes may negatively affect lynx populations by increasing predation on snowshoe hares, and direct killing of lynx (Ripple et al., 2011). The recovery of wolf populations could potentially keep the populations of coyotes and ungulates in check, leading to recovery of plant communities and eventually population growth in snowshoe hares and possibly lynx as well (Ripple et al., 2011). Coyotes also consumed a significant amount of moose (30.10%) and deer (21.36%).

### High Moose Consumption by Wolf in the Dirty Shirt Pack Range

Differences in dietary composition among different wolf pack ranges may reflect prey abundance variations and pack-specific hunting behaviors. Generally, predators, such as wolves, select prey items according to their availability and shift to consuming alternative food sources when the primary food source becomes scarce (Nordberg & Schwarzkopf, 2019; Randa, Cooper, Meserve, & Yunger, 2009). The FOO of moose was the highest (71.43%) in Dirty Shirt, followed by Smackout (38.24%) and Goodman Meadows (24.32%), which implies high moose abundance in the Dirty Shirt range. However, pack-specific hunting ability might also affect prey selection, with larger packs having higher success rates in capturing and killing formidable prey (MacNulty, Tallian, Stahler, & Smith, 2014), such as moose. The Dirty Shirt pack was the largest wolf pack (n = 7 - 13) with confirmed successful breeding pairs during the study period (April in 2015 - May in 2017) (Supplementary Table 2). Larger pack size could make wolves in the Dirty Shirt range more effective at preying on moose. Both pack-specific prey abundance and hunting-ability could affect prey consumption rates. The correlation between moose consumption rate and moose abundance in the Dirty Shirt pack range needs to be further investigated with the inclusion of prey abundance data in this ecosystem.

### Coyote & Ungulate: Ungulate Neonate Predation or Scavenging?

It has been widely assumed that coyotes are not efficient predators on adult deer and are incapable of killing adult moose, but coyotes can occasionally prey on ungulate neonates (<3 months old) during the fawning season (mid-May to mid-June) (Benson & Patterson, 2013; Chitwood et al., 2015; Chitwood, Lashley, Moorman, & Deperno, 2014). Moose calves are vulnerable to coyote predation as well. Coyote pups are born in April, resulting in high lactation demands on females in the spring (Kilgo et al., 2012). This aspect of the coyote life cycle could result in higher pressure on ungulate neonates. The high FOO of moose in the coyote spring diet in the Dirty Shirt range (66.67%) suggests that coyote might predate on moose calves. However, the ungulate neonate predation hypothesis cannot explain the high FOO of deer (60.00%) in the fall coyote diet in the Smackout range, when the deer consumption rate by coyotes in the spring was only 20.83%. Interestingly, moose consumption by wolves was highest in the Dirty Shirt range (71.43%) with no seasonal difference. Deer consumption by wolf was highest in the Smackout range (58.82%), especially in the fall (80.00%). Furthermore, the dietary overlap between these canid species was greatest in the Dirty Shirt range (0.70) in the spring and in the Smackout range in the fall (0.74). Overall, these multiple lines of evidence suggest that coyotes use ungulate carrion subsidies from wolves as a highly-valued food resource. The substantial deer consumption by wolf in the Smackout range in the fall (80%) could be due to severe weather or disease outbreak which might make deer more vulnerable to wolf predation. It could also be due to reduced interspecific competition with other ungulates in this area.

### Consumption of Domestic Animals by Wolves and Coyotes

We detected the DNA of domestic animals in the diets of wolves and coyotes, including pigs, rabbits and domestic cow (Figure 3). Pig DNA was detected in three wolf samples in Goodman Meadows, including one in the fall and two in the spring. Rabbit DNA was detected in two coyote samples, including one in the Dirty Shirt in the fall and the other in the Smackout in the fall. The rabbit DNA was matched to European rabbit (*Oryctolagus cuniculus*). However, this could be from domestic pet rabbits since European rabbits are mainly found on the San Juan Islands in Washington state. In total, we found 18 samples containing cow DNA, including 16 wolf samples and 2 coyote samples. The occurrence of domestic cow DNA in our samples was unlikely to be due to contamination, since none of the negative controls were found to contain cow DNA. Here we only focused on the domestic cow consumption by wolves. Domestic cow was the fourth most frequently occurring prey in the wolf diet, with a FOO of 16.16%. However, it is important to emphasize that the FOO method tends to overestimate the rare prey and underestimate the abundant prey. Indeed, RRA data indicated that the average read proportion of domestic cow in the wolf diet was quite low, only 3.73%. The 16 wolf samples with cow DNA were distributed among three different wolf pack ranges, different years and different seasons. If we assume that 1) all the samples we collected were relatively fresh (based on the high PCR amplification success rate), 2) consumption of any given cow was restricted by the same wolf pack range, in the same season of the same year, and 3) movement of domestic cows across the landscape was limited, we could estimate that there was a minimum of 7 domestic cow individuals involved: two in the Dirty Shirt range (2015 & 2016), two in the Goodman Meadows range in 2015 (fall & spring), and three in the Smackout range (2015, 2016, 2017). Based on Washington Gray Wolf Conservation and Management Annual Reports and gray wolf updates from WDFW, we could only find four reported wolf-cow conflicts that occurred in our study area during the study period, including one incident in Dirty Shirt on Oct 2^nd^, 2016 (confirmed wolf depredation which injured one cow), one incident in Smackout on September 21^st^, 2016 (a confirmed wolf depredation resulting in a dead calf), one incident in Smackout on September 28, 2016 (a probable wolf depredation resulting in a dead calf) and another in Smackout on September 29, 2016 (a confirmed wolf depredation resulting in an injured calf). These four incidents appear to correspond to three samples on the list of Supplementary Table 3, with sample ID 164077 (Dirty Shirt, Fall 2016), 164007 and 164010 (both in Smackout, Fall, 2016). We could not find wolf-predation reports from WDFW that could represent matches of the rest of 16 wolf samples, making it difficult to establish the causes of domestic cow consumption. Though very promising, DNA metabarcoding technology cannot differentiate active predation from scavenging, partial prey consumption or fecal matter consumption. Therefore, caution is needed when interpreting the results from DNA metabarcoding. Conventional methods such as field necropsies and killing-bite wound examination are able to confirm predator identification. Camera trap provides great insights into the feeding behaviors of predators. DNA metabarcoding technology should supplement, not replace conventional methods. Local ranchers, wildlife biologists, and government agencies can work together to examine multiple lines of evidence and combine expertise from each stakeholder to achieve a better understanding of the impact of wolf recovery on the local ecosystem in Washington state.

### Predator-Specific Blocking Primer is Not Necessary with High-Throughput Sequencing Platforms

In dietary studies with fecal DNA, samples contain higher amounts of predator DNA than prey DNA, which can cause PCR amplification being dominated by predator DNA, resulting in low sequencing depth for prey characterization. Predator-specific blocking primers can offer a solution by specifically reducing the amplification of predator DNA. However, with NGS technology becoming faster and cheaper, it might not be necessary to apply predator-specific blocking primer in the diet analyses of carnivores. Indeed, though the majority of sequence reads in our study were from the predator hosts, the amount of prey sequences generated was large enough to characterize the diet profile without the use of blocking primers. Moreover, predator specific blocking oligos can block prey DNA along with the targeted predator (Piñol et al., 2014; Robeson et al., 2017) or cause amplifications to fail altogether (Shehzad et al., 2012), introducing additional bias into the analysis of diet composition. In our study, the use of blocking primers increased the proportion of prey sequences from 26.40% to 65.97%. However, all prey species were detected without adding the blocking primer and chipmunk was not detected when the blocking primer was applied. Given the above, we do not believe that predator-specific blocking primer is cost-effective in the diet analyses of carnivores, and this will likely become even more so as NGS technology continues to become faster and cheaper.

### Recommendations for Future Research

It is important to include negative controls and sequence them along with samples for metabarcoding studies. A low amount of contamination is inevitable with NGS technologies, especially when using universal primers, despite good laboratory practices to minimize contamination risks. We found noticeable contamination in our negative controls with sources from human, striped skunk, wolf, coyotes, moose and deer. Negative controls should always be included to check for potential contamination (De Barba et al., 2013). These controls are often included during steps of DNA extraction and PCR, but they are not always sequenced and may only get checked using gel electrophoresis. Such practice can be misleading as most contaminating sequences cannot be visually detected via gel electrophoresis. By contrast, the series of filtering steps we conducted to remove the impacts of contaminations are very effective and provide valuable framework for contamination control. We also recommend the use of PCR replicates that are sequenced independently as a way to help confirm the presence of taxa in a given sample and further remove false-positives (De Barba et al., 2013; Galan et al., 2018). Pooling PCR replicates prior to sequencing masks the variation among PCR replicates. Therefore, we recommend this multi-replicate approach that the presence of a prey item in a given sample is only confirmed if it occurs in at least 2 replicates.

The key advantage of DNA metabarcoding relative to traditional methods is its high taxonomic resolution. However, this method also has its limitations. Most MOTUs were assigned at the species level except for deer, ground squirrel, red-backed vole, flying squirrel, and chipmunk, which can only be assigned at the genus level. This was likely due to our use of a single short marker (12SV05F/R, ~100 bp). The degree of DNA degradation in fecal samples limits the length of fragments that can be successfully amplified. For this reason, the recommended fragment length is usually in the range of 100-250 bp, which inevitably reduces taxonomic resolution (Pompanon et al., 2011). The multigene approach (Gunther, Knebelsberger, Neumann, Laakmann, & Arbizu, 2018) and mitogenomics approach (Piñol et al., 2014; Tang et al., 2014) have been proposed as the next phase of the current single-marker metabarcoding method. As a PCR-free approach, the mitogenomics approach can alleviate the artifacts caused by PCR bias while expanding single-marker metabarcoding into whole mitochondria metagenomics (Taberlet et al., 2012). The growing mitogenome databases and the continuously decreasing cost of sequencing will eventually make the mitogenomics approach much more affordable and favorable over the single-marker DNA metabarcoding approach.

## Acknowledgements

We thank all the conservation canines and dog handlers who have conducted sample collection for the study, including Heath Smith, Jennifer Hartman, Suzie Marlow, Will Chrisman, Caleb Stanek, Justin Broderick, Casey McCormack, Mairi Poisson, Collette Yee, Rachel Katz, Jake Lammi, Peter Dubyoski, Marlen Richmond and Julianne Ubigau. We thank the Burke Museum of Natural History for providing positive samples during the initial testing phase. We thank Noah Synder-Mackler, India Schneider-Crease, and Sierra Sams for their advice**s** on library prep and MiSeq sequencing. The MiSeq sequencing platform is funded by the Student Technology Fee at the University of Washington. We thank Pierre Taberlet for his advice with experiment design and Susanne Butschkau for her help with bioinformatic data analysis. The study was funded by the Johnson Foundation, the Dawkins Trust and the Maritz Family foundation. Yue Shi was funded by WRF Hall Fellowship from the University of Washington.

## Data Accessibility

After peer review and prior to final publication, the following data will be deposited in Dryad: (i) raw sequence reads (*fastq* format), (ii) filtered abundance table including taxonomic affiliations.

## Author Contributions

YS and SKW conceived the project. YS designed the experiments. YS, YH and ER performed the experiments. YS conducted the analyses and wrote the manuscript. SKW guided analyses and edited the manuscript. All authors declare no conflict of interests.

