## Supplementary for "Prey partitioning between sympatric canid species revealed by DNA metabarcoding"

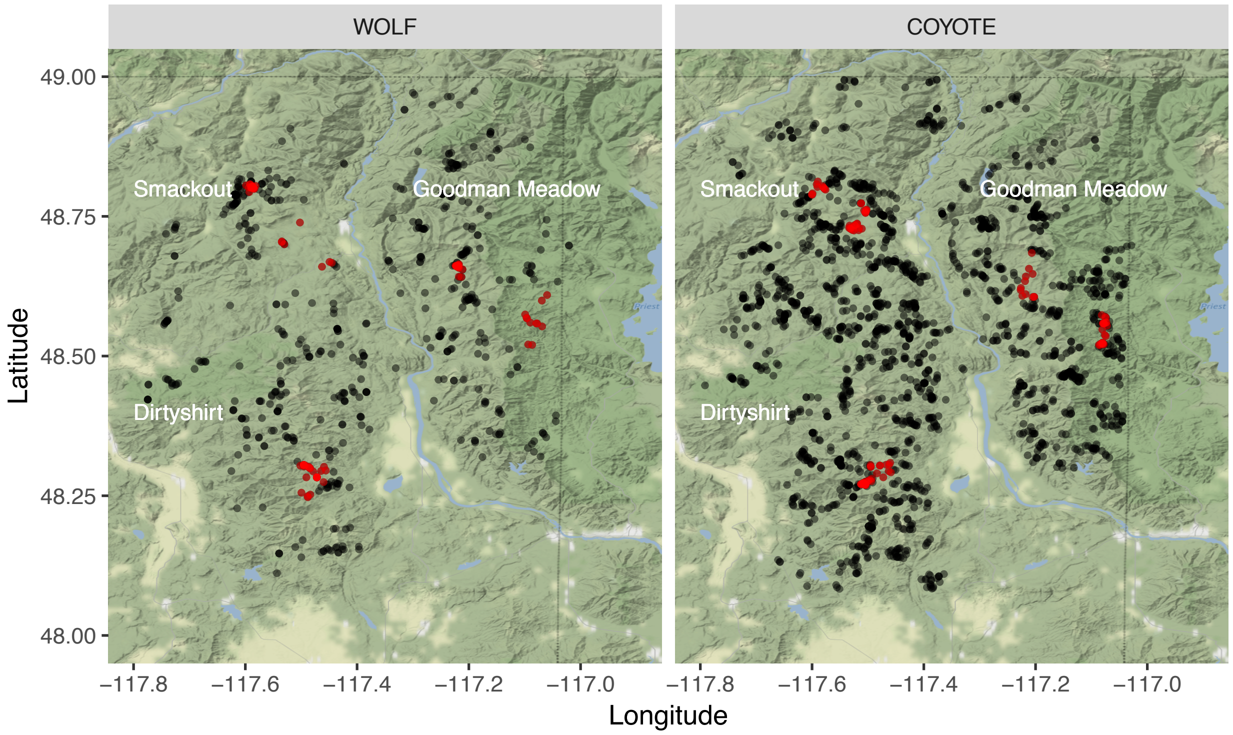


**Supplementary Figure 1** Samples used in this study were selected from an ongoing project with 647 wolf fecal samples, and 1893 coyote fecal samples.

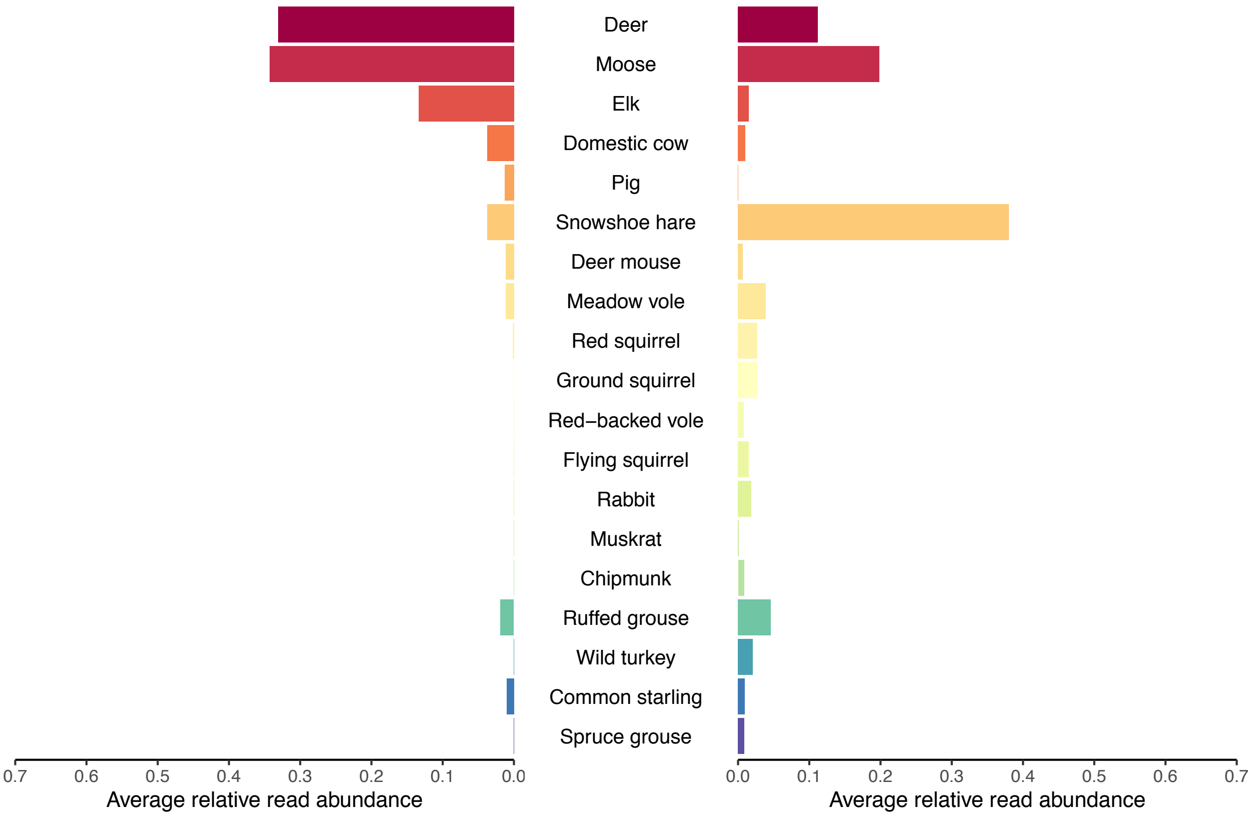


**Supplementary Figure 2** Diet profiles of wolves ($N = 99$) and coyotes ($N = 103$) using the average relative read abundance of 19 prey species.

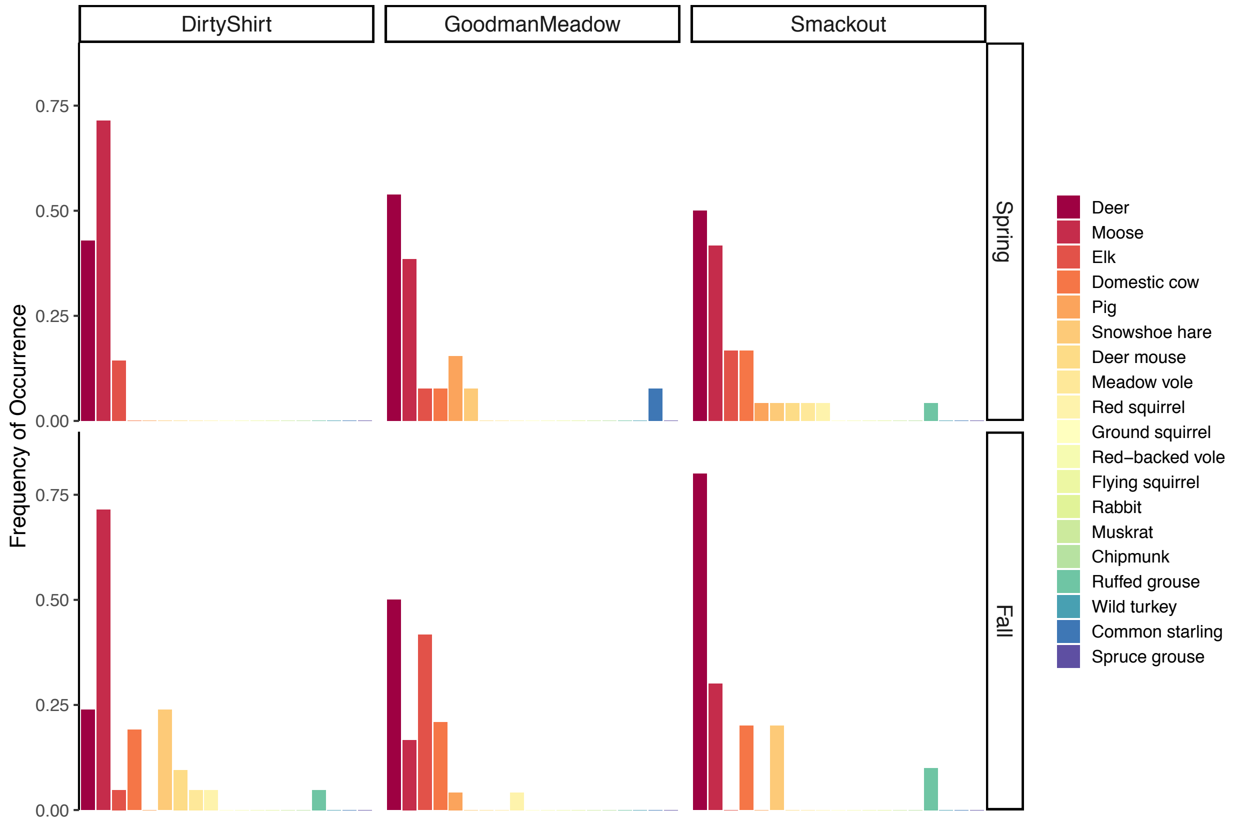


**Supplementary Figure 3** Spatiotemporal variations in the wolf diet profile, using the frequency of occurrence of 19 prey species. The dietary differences among wolf pack ranges were significant ($p$ = 0.001). There was no significant seasonal difference in wolf diet ($p$ = 0.448) nor significant interaction between seasons and wolf pack ranges ($p$ = 0.095).

**Supplementary Table 1** Impacts of the predator-specific blocking primer on the occurrence of difference prey species.

| Prey | Without blocking primer | With blocking primer |
|---|---|---|
| Chipmunk | 1 | 0 |
| Common starling | 2 | 2 |
| Deer | 50 | 60 |
| Deer mouse | 8 | 15 |
| Domestic cow | 13 | 12 |
| Elk | 19 | 22 |
| Flying squirrel | 2 | 3 |
| Ground squirrel | 6 | 6 |
| Meadow vole | 8 | 10 |
| Moose | 60 | 62 |
| Muskrat | 2 | 1 |
| Pig | 4 | 3 |
| Rabbit | 2 | 2 |
| Red squirrel | 5 | 10 |
| Red-backed vole | 3 | 4 |
| Ruffed grouse | 6 | 11 |
| Snowshoe hare | 68 | 65 |
| Spruce grouse | 1 | 1 |
| Wild turkey | 3 | 3 |

**Supplementary Table 2** Wolf pack size from 2015 to 2017 based on Washington Department of Fish and Wildlife Annual Report.

| **Pack** | **2015[1]** | **2016[2]** | **2017[3]** |
|---|---|---|---|
| Dirty Shirt | 8 | 13 | 7 |
| Goodman Meadows | 7 | 7 | 5 |
| Smackout | 8 | 8 | 6 |

Note: [1]: Washington Gray Wolf Conservation and Management 2015 Annual Report (https://wdfw.wa.gov/publications/01793); [2]: Washington Gray Wolf Conservation and Management 2016 Annual Report (https://wdfw.wa.gov/publications/01895); [3]: Washington Gray Wolf Conservation and Management 2017 Annual Repot (https://wdfw.wa.gov/publications/01979);

**Supplementary Table 3** List of samples found to contain domestic cow DNA

| Sample ID | Predator ID | Cow Read Count | Pack Range | Season | Year |
|---|---|---|---|---|---|
| 9183 | COYOTE | 33 | Smackout | Fall | 2015 |
| 847 | COYOTE | 26 | Smackout | Spring | 2015 |
| 164077 | WOLF | 6 | DirtyShirt | Fall | 2016 |
| 9106 | WOLF | 32 | DirtyShirt | Fall | 2015 |
| 9107 | WOLF | 81 | DirtyShirt | Fall | 2015 |
| 9113 | WOLF | 220 | DirtyShirt | Fall | 2015 |
| 7190 | WOLF | 14 | GoodmanMeadow | Fall | 2015 |
| 7191 | WOLF | 528 | GoodmanMeadow | Fall | 2015 |
| 7193 | WOLF | 4 | GoodmanMeadow | Fall | 2015 |
| 7198 | WOLF | 17 | GoodmanMeadow | Fall | 2015 |
| 7204 | WOLF | 1585 | GoodmanMeadow | Fall | 2015 |
| 308 | WOLF | 56 | GoodmanMeadow | Spring | 2015 |
| 164007 | WOLF | 8 | Smackout | Fall | 2016 |
| 164010 | WOLF | 4 | Smackout | Fall | 2016 |
| 170859 | WOLF | 7 | Smackout | Spring | 2017 |
| 829 | WOLF | 127 | Smackout | Spring | 2015 |
| 831 | WOLF | 327 | Smackout | Spring | 2015 |
| 840 | WOLF | 158 | Smackout | Spring | 2015 |
